# mTORC1-Dependent Signaling in Layer 5b Neurons Is Required for Memory Consolidation

**DOI:** 10.64898/2025.12.03.692103

**Authors:** Mik Schutte, Alexander Biermeier, Kinga Konkel, Anna Nasr, Anisha Dayaram, Christian Rosenmund, Robert Sachdev, Matthew E Larkum, Volker Haucke, Marina Mikhaylova

**Author notes:** shared first authorship (Mik Schutte, Alexander Biermeier, Kinga Konkel). shared correspondence.

## Abstract

Memory consolidation relies on activity-dependent neuronal plasticity, particularly in deep-layer cortical neurons that integrate long-range inputs from medial temporal lobe (MTL) structures. In the neocortex, apical tuft dendrites of layer 5b pyramidal neurons receive direct input from the MTL within cortical layer 1, and their activation is critical for associative learning. However, the molecular mechanisms that enable learning-induced plasticity in these neurons remain poorly understood. Here, we demonstrate that activity-dependent, mTORC1-regulated translation in layer 5b neurons contributes to learning and memory. Using in vivo paradigms such as exposure to an enriched environment (EE) and in vitro models of network activation by chemical long-term potentiation (cLTP), combined with cell type-specific translational profiling (RiboTag), we identify candidate mRNAs associated with experience-dependent plasticity. We further show that pharmacological inhibition of canonical mTORC1 signaling during memory-relevant tasks abrogated memory consolidation. Our findings identify the mTORC1 pathway as a central regulator of learning-induced protein translation and cortical plasticity.

## Introduction

Systems memory consolidation describes the gradual reorganization of hippocampal-dependent memories into distributed neocortical networks for long-term storage (Squire, 1986; Frankland and Bontempi, 2005; Kitamura et al., 2017). Recent studies identify neocortical layer 1 as a principal site of long-term plasticity and learning (Cichon and Gan, 2015; Abs et al., 2018; Williams and Holtmaat, 2019; Pardi et al., 2020). Serving as a gateway for long-range inputs from the medial temporal lobe and higher-order thalamus (Rubio-Garrido et al., 2009; Yang et al., 2016; Shin et al., 2021) these inputs converge specifically onto layer 1 that contains the apical tuft dendrites of layer 5b pyramidal neurons, suggesting these dendrites are a crucial locus for memory stabilization. However, the molecular mechanisms that convert these transient synaptic inputs into lasting structural changes remain poorly defined.

Long-term memory formation requires *de novo* protein synthesis (Davis and Squire, 1984; Costa-Mattioli et al., 2009), implicating translational control as a vital step in consolidation. Recent transcriptomic profiling has confirmed that neocortical layer 1 is a site of active protein synthesis with a rich repertoire of localized synaptic transcripts (Spanò et al., 2025). The mechanistic target of rapamycin complex 1 (mTORC1) signaling pathway has emerged as a central integrator of neuronal activity and translation. While inhibition of mTORC1 disrupts long-lasting synaptic plasticity and long-term memory formation across various species, the specific spatiotemporal engagement of this pathway in neocortical circuits during learning remains unresolved.

Here, we test the hypothesis that activity-dependent, mTORC1-regulated signaling in layer 5b neurons supports cortical memory consolidation. We compared a layer 1-dependent cortical microstimulation learning paradigm (Doron et al., 2020) with enriched environment exposure (Rampon et al., 2000; Olson et al., 2006) and *in vitro* models of network activation (Villers et al., 2014). Using cell-type-specific RiboTag profiling of translating ribosomes we identified a coordinated translational program associated with plasticity and mTORC1-dependent signalling emerged as a promising candidate pathway. Furthermore, we demonstrate that pharmacological inhibition of canonical mTORC1 signaling during the consolidation phase disrupts memory formation. Altogether, our findings identify mTORC1 signaling in layer 5b neurons as a critical checkpoint for stabilizing cortical memory, coincident with the upregulation of a cell-type-specific translational program.

## Materials & Methods

### Animals

All experiments involving the use of animals were conducted in accordance with the European Communities Council Directive (2010/63/BJ) and the Animal Welfare Law of the Federal Republic of Germany (Tierschutzgesetz der Bundesrepublik Deutschland, TierSchG) approved by the local authorities of the city-state Berlin (Behörde für Gesundheit und Verbraucherschutz, Fachbereich Veterinärwesen) and the animal care committee of the Humboldt University Berlin. Animals used for the study were bred and housed in the animal facility of the HU-Berlin (T-CH 0011/20, G 0146/21). Animals were housed under a reversed 12-hour light/dark cycle (lights on at 21:00, off at 09:00), and all behavioral training was conducted during the dark phase.

### Enriched Environment Experiments

Mice were placed either in an enriched environment or in a standard home cage. Exploratory activity, defined as the percentage of time spent exploring, was measured over a 120-min session to determine periods of highest activity (Supplementary Fig. 2a). To quantify activity, we developed an automated tracking and analysis tool, ActiveMouse (https://github.com/Lilli-K2/ActiveMouse-SFB1315), which integrates the DeepLabCut-Live framework (Kane et al., 2020) with custom in-house code to monitor and analyse mouse activity in real time, without requiring researcher presence. The software generates (I) a heat map showing frequently visited areas (Supplementary Fig. 2b), (II) a tracking plot indicating total tracking duration (Supplementary Fig. 2c), and (III) a summary plot of total distance travelled and mean movement speed (Supplementary Fig. 2d). All results are automatically produced at the end of the session, enabling immediate verification that each mouse met the predefined activity criteria required for inclusion. This approach allows exclusion of inactive animals prior to sacrifice, thereby reducing the number of mice used in accordance with the 3R principles.

For spatial transcriptomic experiments four female C57BL/6J wild type (WT) mice, 5 weeks old were used. The animals were housed alone for 4 hours prior to experiment and were on a reverse day/night schedule three days prior. Two sets of experiments took place, one in the early morning and one in mid-morning. In both experiments one animal was placed in a novel enriched environment whilst the second animal was placed in the standard cage for 30 minutes. Experiments were filmed using a night vision camera (Basler acA4096-30um, Song Pregius) to ensure animals explored the cages of the enriched environment. Following the experiment both animals were euthanized with isoflurane with subsequent cerebral dislocation.

### Spatial Transcriptomics Tissue Preparation

The brains from both the enriched environment and animals kept in standard environment were subsequently removed and placed directly in a cryomold and embedded with Optimal Cutting Temperature solution (OCT, TissueTek), and placed on dry ice. OCT blocks were kept at –80 °C for sectioning. Cryosectioning was performed for each of the four animals on different days using a Leica CM3050S cryostat at a temperature setting of –20 °C for the blade and –10 °C for the specimen head with the humidity between 36-64%. 10µm coronal tissue sections were made from both right and left hemispheres of the brain focusing on the S1 barrel region. Each right and left hemisphere from each hemisphere was placed on the Visium slide within the 6.5 x 6.5 mm square. Slides were kept frozen during sectioning. Slides were fixed with methanol at –20 °C and stained with hematoxylin and eosin (H/E) according to the standard 10X Visium protocol.

Imaging of the H/E staining was captured using the Nikon Eclipse Ti2 microscope (AMBIO facility, Charité). Initially, the Visium imaging test slides were imaged using both brightfield and fluorescence to select optimal tissue lysis time at 30 minutes (see Visium protocol). Following tissue optimization, Visium spatial gene expression slides were imaged using brightfield in 24bit colour, and tiles were subsequently stitched together and saved as TIF and nd2 files.

### Spatial Transcriptomics Library Preparation and Sequencing

Following the slide imaging cDNA and library building was completed for both slides following the standard 10X Visium Spatial Transcriptomics protocol. Libraries were pooled and quantified based with both the Agilent Bioanalyzer using high sensitivity chip and reagents (Agilent Technologies, USA) and the Agilent qPCR NGS Library Quantification kit (Agilent Technologies, USA).

Sequencing was carried out at the BIH/MDC Genomics facility (Berlin, Germany) on the Illumina NovSeq 6000 platform using one lane of an S4 V1.5 flowcell with paired-end reads.

### Spatial Transcriptomics Bioinformatic Analysis

For spatial transcriptomic data file processing, UMAP analysis and spatial feature plotting was analyzed using Seurat package, v5.2.1. For cell deconvolution analysis of spatial transcriptomic data, the same workflow was implemented for single-cell transcriptomics reference data from the Allen Mouse Brain Atlas (Lein et al., 2007). This was then used to anchor our own data with the brain regions as well as the cell types. Robust Decomposition of cell type mixtures (Cable et al., 2022) analysis was carried out with the Spacexr package, v2.2.1. Differential gene expression was based on the DESeq2 version 1.46.0 (Love et al., 2014). All spatial transcriptomics analyses and figures were produced in R, v4.4.2.

### Primary cortico hippocampal-cultures

Primary neuronal cultures were prepared from wild-type C57BL/6J mice (Charles River). Cortico-hippocampal neurons were isolated from neonatal (P0) mouse brains. Cortices and hippocampi were dissected and dissociated with trypsin (1 mg/mL; Sigma-Aldrich) and DNase. Cells (70,000–100,000/cm²) were plated on poly-L-lysine–coated coverslips or plates in plating medium [MEM supplemented with 0.5% glucose, 0.02% NaHCO₃, 0.01% transferrin, 10% FCS, 2 mM L-glutamine, insulin (25 μg/mL), and 1% penicillin–streptomycin]. On DIV1–2, medium was partially replaced with growth medium containing AraC (2 µM) to reduce glial proliferation. From DIV7 onward, cultures were maintained by partial medium exchange with BrainPhys (STEMCELL Technologies, #05790) supplemented with NeuroCult SM1 (STEMCELL Technologies, #05711) and 1% penicillin–streptomycin.

### Neuronal activation with cLTP

For RiboTag experiment, neurons were stimulated using a chemical long-term potentiation protocol (cLTP) in artificial cerebrospinal fluid solution (ACSF) [25 mM HEPES, 125 mM NaCl, 2.5 mM KCl, 2 mM CaCl2 and 33 mM glucose (pH 7.4)]. Control cells were treated with ACSF containing 1 mM MgCl_2_ to block NMDA receptors and stimulated ones treated with a buffer without MgCl_2_ to relieve the NMDA receptors block. Neurons were washed once with appropriate ACSF solution and then stimulated for 15 min with a mixture of glycine (100 µM; Roth, 3908.2), strychnine hydrochloride (1 µM; Sigma-Aldrich, S8753) and bicuculline (25 µM; Tocris, 2503) in ACSF without MgCl_2_. After 15 min, cLTP solution was replaced with fresh ACSF and incubated for another 45 min.

### Translating Ribosome Affinity Purification (TRAP) of RPL22-3xHA-mCherry expressing primary neurons and eGFP-L10a mouse line

A lentiviral construct expressing RPL22-3xHA-RFP under the human Synapsin1 promoter was generated using Gibson assembly. The lentiviral backbone (gift of C. Rosenmund) and the RPL22-3xHA-mCherry sequence (Addgene #170320; H. Phatnani) served as templates. Lentiviral particles were produced in HEK293T cells supplemented with nonessential amino acids and sodium pyruvate, using calcium phosphate transfection of the transfer vector together with psPAX2 and pMD2.G helper plasmids. Viral supernatants were harvested at 48 h and 72 h, clarified by filtration, concentrated using 100 kDa centrifugal filters, and stored at −80 °C.

For *in vitro* TRAP, primary cortico-hippocampal neurons were transduced with lentiviral particles expressing RPL22 tagged with HA and mCherry at DIV2. After 1h of cLTP treatment, cells were washed with ice-cold TBS and resuspended in ice-cold polysome lysis buffer [50 mM Tris-HCl (pH 7.4), 100 mM KCl, 5 mM MgCl2, 1 mM dithiothreitol (DTT), 100 µg/ml cycloheximide (CHX), 200 U/mL RNase (Promega), 1 mg/ml heparin, Pierce™ Protease Inhibitor Mini Tablets, EDTA-free (A32955; Thermo Scientific), phosphatase inhibitor cocktail 2&3 (P5726 & P0044; Sigma-Aldrich), 1% NP-40]. Cells were lysed by constant rotation for 20 min at 4°C and cleared by centrifugation. Protein concentration was measured using Pierce™ BCA Protein Assay Kit (23227; Thermo Scientific). Equal amounts of protein lysate were loaded onto Pierce™ Anti-HA Magnetic Beads (88836; Thermo Scientific) that were previously washed twice with polysome lysis buffer and incubated overnight at 4°C using end-over-end mixing. After incubation, flow-through samples were transferred to a new tube, combined with Laemmli Sample Buffer [final (1 x) concentration: 31.5 mM 2-Amino-2-(hydroxymethyl)propane-1,3-diol (Tris), 1% (w/v) sodium dodecyl sulfate (SDS), 10% (v/v) glycerol, 0.001% (w/v) 3,3-Bis(3,5-dibromo-4-hydroxyphenyl)–2,1λ6-benzoxathiole-1,1 (3H)-dione (bromophenol blue), 5% (v/v) 2-mercaptoethanol; pH 6.8] and boiled at 95°C for 5 min. Beads were washed three times with high salt polysome buffer (polysome lysis buffer with 300 mM KCl) with end-over-end mixing at 4°C. After removing all washing buffer, beads were resuspended in RLT buffer from RNeasy extraction kit (74104; Qiagen) supplemented with 0.01% β-mercaptoethanol.

For the *in vivo* TRAP, mouse strain Glt25d2-EGFP/L10a (JAX stock no. 030257 (Doyle et al., 2008)) was used for the purification of cortical translating mRNA from layer 5. Adult male mice (> 9 weeks old; n = 16 total) were used. Eight animals were used for each condition (enriched environment, standard environment), and tissue from four mice was pooled to create one biological sample (n = 2 pooled biological samples per condition). The mice were decapitated after exposure for 30 or 90 min to enriched environment experiments, the meninges were removed and the cortical tissue was manually dissociated and immediately frozen away in liquid nitrogen. The pooled tissue was then homogenised on ice using a motor-driven Teflon glass homogeniser in ice-cold supplemented homogenisation buffer [50 mM TRIS (pH 7.4) (Roth, 4855.2), 100 mM KCl (Roth, 6781.1), 12 mM MgCl_2_ (Sigma, M4880-100g), 1 mg/ml heparin (Sigma, H3393), 1x protease inhibitor (Roche, 11484100), 1000 µg/ml cycloheximide (Sigma, c7698-1g), RNase inhibitor (Promega, N2515), 1 mM DTT (Roth, 6908.2)]. The homogenised mixture was left on ice for 10 min until the foam settled and then for 10 min at 2000 x g, 4 °C, to separate the cell nuclei from the lysate. Consequently, 10% IGEPAL CA-630 (Sigma-Aldrich, 18896) was added to the supernatant to obtain a final concentration of 1 % and inverted until the lysate appeared clear. After incubation for 5 min on ice, the lysate was centrifuged for 10 min at 13,000 x g to pellet insoluble material. Rabbit anti-GFP-coated (see beads preparation) protein A and G magnetic beads were added to the supernatant and the mixture was incubated at 4 °C overnight. The beads were then collected using a magnetic rack on ice and washed 3 times with a high salt washing buffer [300mM KCl, 10 % IGEPAL, 50 mM TRIS [ph 7.4], 12 mM MgCl2, 1mM DTT, 100 µg/ml cycloheximide] and resuspended in Trizol-LS reagent (Invitrogen; 15596026) to isolate the bound mRNA from the ribosomes. After extraction, the RNA was precipitated with 95-100% ethanol and further purified using the Direct-zol Microprep Kit (Zymo Reasearch, R2060) following the instructions. The isolated RNA was diluted nuclease-free water and stored at –80° C and further used for mRNA sequencing. RNA concentration and purity was measured using NanoDrop 2000(Thermo Fisher Scientific, Wilmington, DE). RNA integrity was assessed using the RNA Nano 6000 Assay Kit of the Agilent Bioanalyzer 2100 system (Agilent Technologies, CA, USA)

For bead preparation, magnetic beads protein A and protein (BioradSureBeads Cat#161-4013 and Cat#161-4023,) were mixed in a 1:1 ratio and washed twice with a homogenisation buffer [50 mM TRIS [pH 7.4], 100 mM KCl, 12 mM MgCl2]. The beads were then incubated with a homogenisation buffer and 0.2% chicken egg white (Sigma, A5503) and anti-GFP (abcam, ab6556) in rotation at 4 °C overnight. On the following day, the beads were washed once with a supplemented homogenisation buffer and resuspended in the original volume.

### Immunohistochemistry and Immunocytochemistry

For immunohistochemical staining of target proteins we used Ai9xSim-Cre mice (n=6; 1 male, 5 females; age=9.8 ±0.5 weeks) and C57BL/6J mice (n=6, 6 males; age=9.1 ±0.11 weeks) mice. After the behavioural test, they were anaesthetised with Ketamine/Xylazine (65mg/kg;10mg/kg, i.p.) and transcardially perfused with 1X phosphate buffer saline [PBS 8.05 mM Na_2_HPO_4_ (Roth, P030.1), 1.95 mM KH_2_PO_4_ (Roth, P030.1), 150 mM NaCl (Roth, P029.3) [pH 7.4]] followed by 4% paraformaldehyde PFA (Roth, 335.1) in PBS. The brains were removed and postfixed in 4 % PFA for at least 24 h and then transferred to 30% sucrose (VWR, 27480294) solution in PBS for 48-72 h until the brains had settled. The brains were then frozen on dry ice in Tissue tek® O.C.T. compound. Coronal brain sections were cut into 40 µm thick slices using a cryostat and then stored at 4 °C in PBS. The brain sections were incubated for 30 min at room temperature (RT) in blocking buffer [PBS, 0.3% Triton-X 100 (Sigma-Aldrich, X100), 10% goat serum (Gibco; 16210064)]. This was followed by incubation with primary antibody diluted in blocking buffer for 36 h at 4 °C on a shaker. The brains were then extensively washed in PBS for 1 h with multiple changes of wash buffer. Subsequently, the sections were incubated in PBS containing 0.2 % bovine serum albumin BSA (Roth, 8076.2) for 1 h. The sections were then incubated with fluorescently labelled secondary antibodies for 2h. After washing with PBS, the sections were mounted on slides with Mowiol (Merck, Darmstadt, Germany 81381).

For immunocytochemistry, neurons were fixed at DIV14 with 4% PFA, 4% sucrose in PBS for 15 min at RT, washed, and blocked in 0.2% Triton X-100, 10% normal goat serum for 1 h. Cells were incubated with primary antibodies in a permeabilization buffer overnight at 4 °C, washed, and incubated with secondary antibodies for 1 h at RT. After final PBS washes including DAPI, coverslips were rinsed, mounted with Immu-mount, and dried overnight protected from light.

### RNA sequencing

For *in vivo* samples: Sequencing libraries were generated using NEBNext UltraTM RNA Library Prep Kit for Illumina (NEB, USA) following manufacturer’s recommendations and index codes were added to attribute sequences to each sample. The qualified library was sequenced on the Illumina Novaseq X platform (Illumina, San Diego, USA) with paired-end 150 bp (PE150) mode. Library construction and sequencing were performed at Biomarker Technologies (BMKGENE) GmbH.

For *in vitro* samples: Isolated RNA was extracted and sequenced using the BGI DNBSEQ platform (http://biosys.bgi.com) to generate both mRNA and lncRNA libraries, with paired-end 100 bp reads and an average yield of approximately 23 million clean reads per sample. Sequencing reads were aligned to the *Mus musculus* genome (Mus_musculus_10090.UCSC.mm39.v2201) using HISAT2 alignment software, v2.0.4 (Kim et al., 2015). Gene expression levels were quantified and normalized to FPKM using RSEM software, v1.2.18 (Li and Dewey, 2011). Differential expression analysis between control and cLTP groups was performed with the DESeq2 package, v1.4.5 (Love et al., 2014).

### RNAseq data analysis

Principal component analysis and visualisation was performed using the Dr. Tom analysis software provided by BGI. Gene ontology (GO) and gene set enrichment analyses (GSEA) were performed using R package clusterProfiler, v4.16.0 (Wu et al., 2021) and <u>org.Mm.eg.db</u>, v3.21.0 (Carlson, 2017) for mouse genes annotation. For GSEA comparison, GO Biological Process (BP) ontologies were used as the sets for enrichment and significance was set to p-value of <0.05. Volcano plots were generated using ggplot2 (Wickham, 2016) and Venn Diagrams using VennDiagram (v1.7.3) R package (Hanbo Chen, 2011).

### Western Blot (WB)

The samples for immunoblotting were diluted using 4 x SDS loading dye [250 mM Tris-HCl (Roth, 9090.4), 8% [w:v] SDS (Roth, CN30.2), 40% [v:v] glycerol (Roth, 3783.1), 20% [v:v] β-mercaptoethanol (Sigma, M3148), 0.008% bromophenol blue (Roth, A512.2), [pH 6.8]] was added and samples were incubated for 5 min at 95°C. SDS-PAGE was performed using 4-20% Mini-PROTEAN TGX Stain-free Protein Gels (Bio-Rad, 4568095) or self-made Bis(2-hydroxyethyl)amino-tris(hydroxymethyl)methane (Bis-Tris; 250 mM) based 8–15% polyacrylamide (Rotiphorese Gel 30; Carl Roth) gradient gels and run (80–130 V; 90 min) in MOPS running buffer (1 M MOPS, 1M Tris base, 69.3 mM SDS, 20.5 mM EDTA) using Mini-PROTEAN Tetra Vertical Electrophoresis Cells (Bio-Rad, Cat#1658004).

For WB the proteins were transferred onto an activated polyvinylidene fluoride (PVDF) membrane (Bio-Rad, 1620177) in transfer buffer (Bio-Rad, 10026938) using a semi-dry blotting system (Bio-Rad). Blocking was done with 5% [w:v] milk powder (Sucofin) in TBS containing Tween20, TBS-T, 20 mM Tris (Roth, 4855.2), 150 mM NaCl (Roth, P029.3), 0.1% [v:v] Tween20 (Roth, 9139.1), [pH 7.4] at RT for 60 min or transferred to nitrocellulose membrane, blocked with Intercept (TBS) Protein-Free Blocking Buffer (LICORbio; 927-60001) for 30 min at RT. Primary antibody incubation was done at 4°C overnight.

All antibody dilutions were made in TBS containing sodium azide TBS-A, 20 mM Tris (Roth, 4855.2), 150 mM NaCl (Roth, P029.3), 0.02% [w:v] sodium azide (Roth, 4221.1), pH [7.4]. Secondary antibodies used were goat anti-mouse-HRP (1:10 000 dilution in blocking buffer, (Dianova, 115-035-146) and goat anti-rabbit-HRP (1:10 000 dilution in blocking buffer (Dianova, 111-035-144). The blots were developed using Pico ECL (Thermo, 34577) on a Bio-Rad ChemiDoc MP device and by Image Studio Lite (Version 5.2.5).

### Microscopy

Confocal imaging was performed with a Nikon Eclipse Ti-E Visitron SpinningDisk confocal microscope controlled by VisiView software (Visitron Systems). The samples were imaged using a 10 x objective (Nikon, 10x Plan Apo Air, NA 0.45) and a 60x objective (Nikon, 60x Plan Apo λ oil, NA 1.40) resulting in a pixel size of 650 nm for 10x and 108.3 nm for 60x with 405, 488, 561 and 640 nm excitation laser. Lasers were coupled to a CSU-X1 spinning disk (Yokogawa) unit via a single-mode fibre. Emission light was collected through a filter wheel with appropriate filters (Chroma ET460/50m, Chroma ET525/50m, ET609/54m and Chroma ET700/75m). Z*-*stacks were acquired with a pco.edge 4.2 bi sCMOS camera (Excelitas PCO GmbH) with 350 nm step size in 16-bit depth.

### Habituation

Mice were gradually habituated to handling across three consecutive days, beginning with 5 minutes on the first day and increasing in 5-minute increments until animals remained calmly on the experimenter’s hand for at least 45 minutes. Two days following headpost implantation, animals were introduced to brief (5-minute) head-fixation. Head-fixation training continued with increments of 10 minutes until the animal was comfortable in the setup for 60 minutes. From the third day onward, mice were placed on a water restriction schedule (1 mL/day) and trained to lick from a piezo-equipped spout.

### Surgical Procedures

To ensure stable head-fixation and precise electrode targeting, mice were implanted with a custom-designed stainless steel headpost. Surgical procedures were performed under isoflurane anesthesia (induction at 3–4%, maintenance at 1.5–2%) with continuous monitoring of respiration and reflexes. Prior to incision, 40 μL of lidocaine (10mg/mL) was administered subcutaneously to the scalp for local analgesia.

The scalp was shaved, and the animal was positioned in a stereotaxic frame with ear bars and a bite bar to ensure head stability. After removing the scalp and the periosteum, the skull surface was briefly treated with a drop of hydrogen peroxide (0.3% in PBS) to aid in tissue cleaning and whitening. After 30 seconds, the area was thoroughly rinsed with sterile PBS (1X) and cleaned with 70% ethanol.

A thin layer of dental adhesive (Optibond, Kerr) was applied to the skull and allowed to settle for 30 seconds before being polymerized under UV light for 60 seconds. Target coordinates for the craniotomy (–1.3AP –3.3ML) were then marked, and the left hemisphere was coated with Rely-X composite resin (manufacturer) to support headpost anchoring. The metal headpost was affixed atop the prepared skull surface using dental cement (Tetric-Evoflow) and allowed to cure completely. Finally, the remainders of the scalp were adhered to the skull.

Two days prior to the onset of microstimulation training, a 1.5 mm craniotomy was performed above the right primary somatosensory cortex (S1; AP: –1.3 mm, ML: –3.3 mm from Bregma). The dura mater was left intact. A custom-made recording chamber was adhered around the craniotomy to allow chronic access, and the craniotomy was sealed with silicone elastomer (Kwik-Cast, World Precision Instruments) to preserve tissue integrity between sessions.

Post-operative analgesia was provided via subcutaneous administration of Carprofen and Buprenorphine, followed by one more day of Carprofen. Animals were monitored during recovery and given at least 24 hours before behavioral procedures resumed

### Microstimulation detection task

Animals were trained to report intracortical microstimulation (µStim) with a tongue-lick response. Stimulation consisted of a 200 ms train of cathodal current pulses (40 pulses; 200 Hz; 0.3 ms duration) at 40 µA delivered via a tungsten microelectrode (MicroProbes) inserted approximately 750 μm below the pial surface in the right primary somatosensory cortex.

On each of the four days of training animals underwent 150 trials divided into 60% µStim and 40% catch trials. All µStim trials were reward-contingent: animals were only rewarded if they licked within a designated response window of 300–1700 ms after stimulus onset. Catch trials lacked a microstimulus, but quantified licks during the same response window to measure the selectivity of behavior and improve trial randomness.

To prevent anticipatory licking, the intertrial interval (ITI) was randomly set between 3 and 8 seconds. In addition, licking during the ITI resulted in the incursion of a new ITI that had to run out before a new trial occurred. This aimed to encourage selective licking after a mStim to enhance the reliability of our behavioural measurements.

Stimulation was delivered at a constant intensity of 40 μA throughout all training sessions. Animals continued training until they completed 150 trials or were in the setup for more than 1.5 hours. An animal was considered expert once it achieved a predefined behavioral performance threshold (see Behavioral Criteria). Training was carried out across 4 consecutive days for animals in the injection experiments, whereas animals used for immunohistochemistry experiments underwent only 2 experimental days.

### Behavioural quantification

The performance of animals during the microstimulation detection task was quantified using the sensitivity index (d’). This index, based on signal detection theory, reflects both the animal’s response to the microstimulus, as well as the animal’s bias to lick (Carandini and Churchland, 2013). The licking bias is captured by the number of False Alarms, i.e. when the animal reported the stimulus when it was not present (Catch Trials). Thus, the sensitivity index compares false alarm rates to hit rates within the same session. In order to calculate d’ we use the following formula d’ = z(HIT) – z(FP) where z is the normal inverse of the cumulative distribution function of the normalized hit and false positive rates. To avoid infinite d’ values we substitute the absolute rate with (1-1/2N) or (1/2N) where N is the number of microstimulation trials. Higher values of d’ indicate a better sensitivity to the microstimulation.

### Pharmacological manipulation of mTORC1 activity

Rapamycin was administered intracortically immediately after each µStim training session to inhibit mTORC1. Mice received either rapamycin (2 mM in DMSO, 200 nL) or saline. Injections were delivered ∼15 minutes after training at 250 µm depth using a glass micropipette (20 nL/s). The pipette was left in place briefly to minimise backflow. Injections were performed once per day for three consecutive training days.

Antibodies

Primary antibodies:

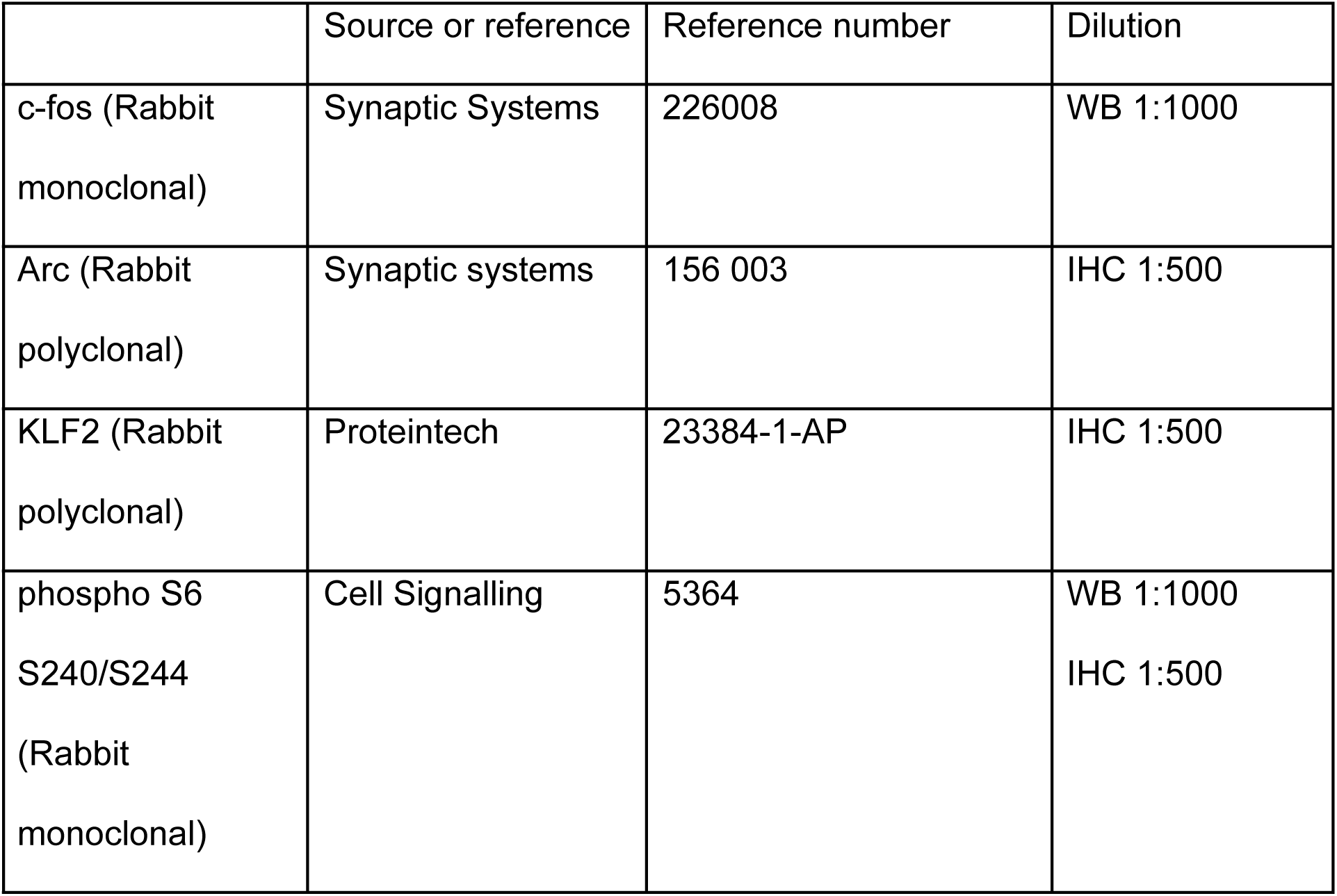
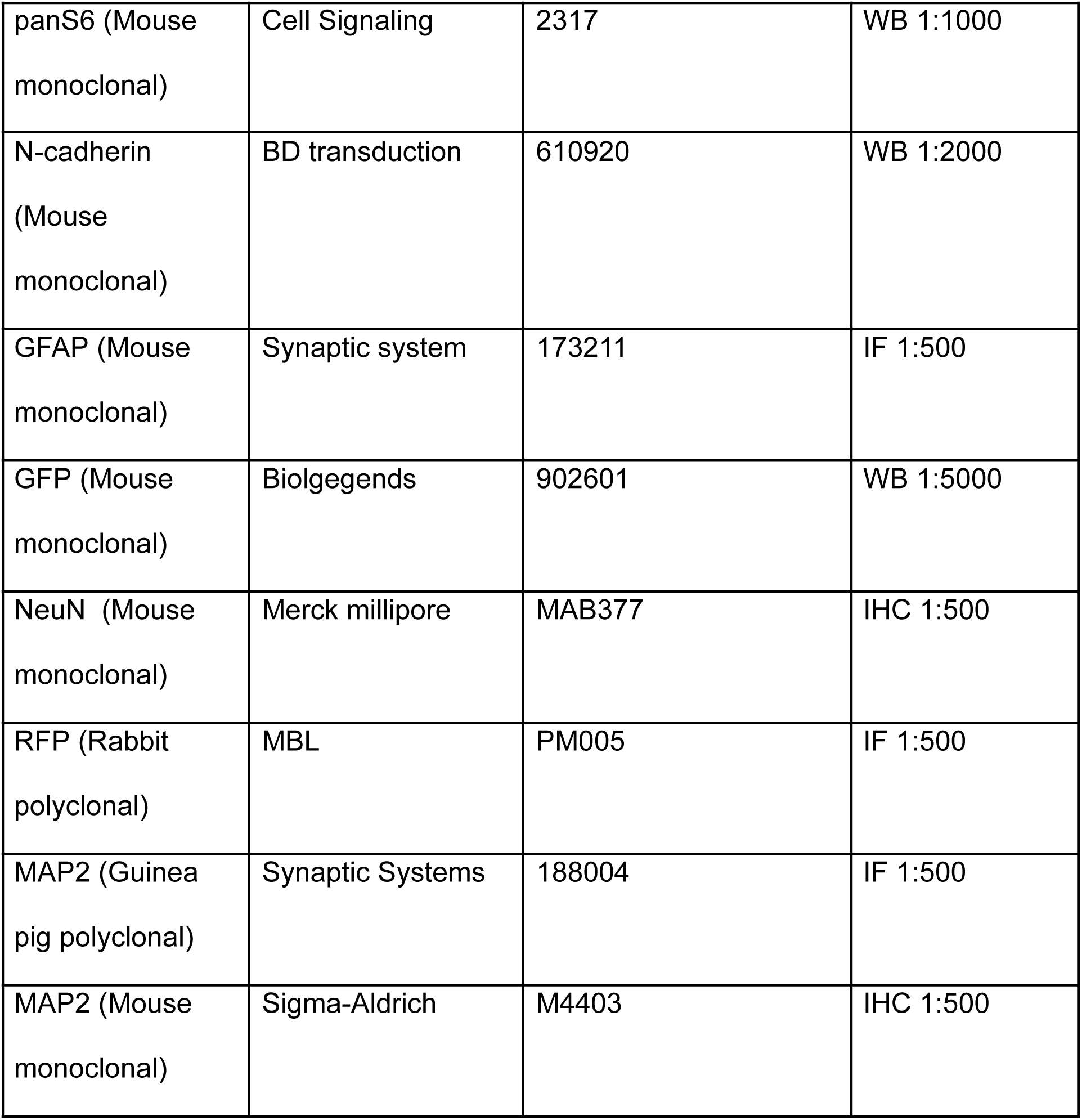

Secondary antibodies:

|  | Source or reference | Reference number | Dilution |
| --- | --- | --- | --- |
| Anti-rat Alexa 488 (donkey polyclonal) | Thermo Fisher Scientific | A21208 | IF 1:400<br>IHC: 1:500 |
| Anti-rabbit Alexa<br>568 (goat<br>polyclonal) | Thermo<br>Fisher<br>Scientific | A11011 | IF 1:400<br>IHC: 1:500 |
| Anti-mouse<br>Alexa 488 (goat<br>polyclonal) | Thermo<br>Fisher<br>Scientific | A11001 | IF 1:400<br>IHC: 1:500 |
| Anti-guinea pig<br>Alexa 647 (goat<br>polyclonal) | Thermo<br>Fisher<br>Scientific | A21450 | IF 1:400<br>IHC: 1:500 |
| Anti-rabbit Alexa<br>647 (goat<br>polyclonal) | Thermo<br>Fisher<br>Scientific | A21245 | IF 1:400<br>IHC: 1:500 |
| Anti-mouse<br>IRDye 680RD<br>(goat polyclonal) | LI-COR<br>Biosciences | Cat# 925–68070;<br>RRID:AB_265112<br>8 | WB 1:5000 |
| Anti-rabbit<br>IRDye 680RD<br>(goat polyclonal) | LI-COR<br>Biosciences | Cat# 926–68071;<br>RRID:AB_10956<br>166 | WB 1:5000 |
| Anti-mouse IgG,<br>IRDye 800CW<br>(goat polyclonal) | LI-COR<br>Biosciences | Cat# 926–32210;<br>RRID:AB_62184<br>2 | WB 1:10000 |
| Anti-rabbit IgG,<br>IRDye 800CW<br>(goat polyclonal) | LI-COR<br>Biosciences | Cat# 926–32211;<br>RRID:AB_62184<br>3 | WB 1:10000 |

## Results

### Activity-dependent translation in cortical neurons engages the mTORC1 pathway

To test the hypothesis that neuronal activity triggers mTORC1 activation and downstream protein translation during memory consolidation, we first sought to identify mRNA targets that undergo activity-regulated protein synthesis. We combined a protocol for the induction of neuronal activity (often referred to as chemical LTP, cLTP) (Fig. 1A, Fig. S1) with RiboTag technology and RNA sequencing in neurons in culture (Fig. 1B). We verified the neuron-specific expression of a triple HA-tagged variant of the large ribosomal subunit eL22 (RPL22) in lentivirally transduced cortico-hippocampal neurons (Fig. 1C). Moreover, we confirmed that neuronal stimulation by our cLTP protocol led to the activation of mTORC1 signaling monitored by the phosphorylation of its downstream substrate ribosomal S6 protein and to the induction of immediate-early gene (IEG) expression as exemplified by c-Fos (Fig. 1D). We then subjected lysates of cortico-hippocampal neurons to cLTP induction and corresponding mock controls to RiboTag affinity chromatography and quantitative analysis by RNA sequencing to identify newly translated proteins. As expected, we found cLTP-induced expression of IEGs such as c-Fos, Fosl, c-Jun, and Arc among others (Fig. 1E). Gene Ontology (GO) enrichment analysis of differentially translated proteins showed that neuronal stimulation induces the translation of proteins involved in RNA splicing, ribonucleoprotein complex biogenesis, protein phosphorylation (e.g. cell signaling), and metabolism (Fig. 1F). Moreover, Gene set enrichment analysis (GSEA), a method that uses gene ranking to detect enrichment of entire pathways, showed that cLTP-induced translation may affect pathways related to angiogenesis, ERK1/2 signaling, as well as learning and cognition (Fig. 1G).

**Figure 1.**
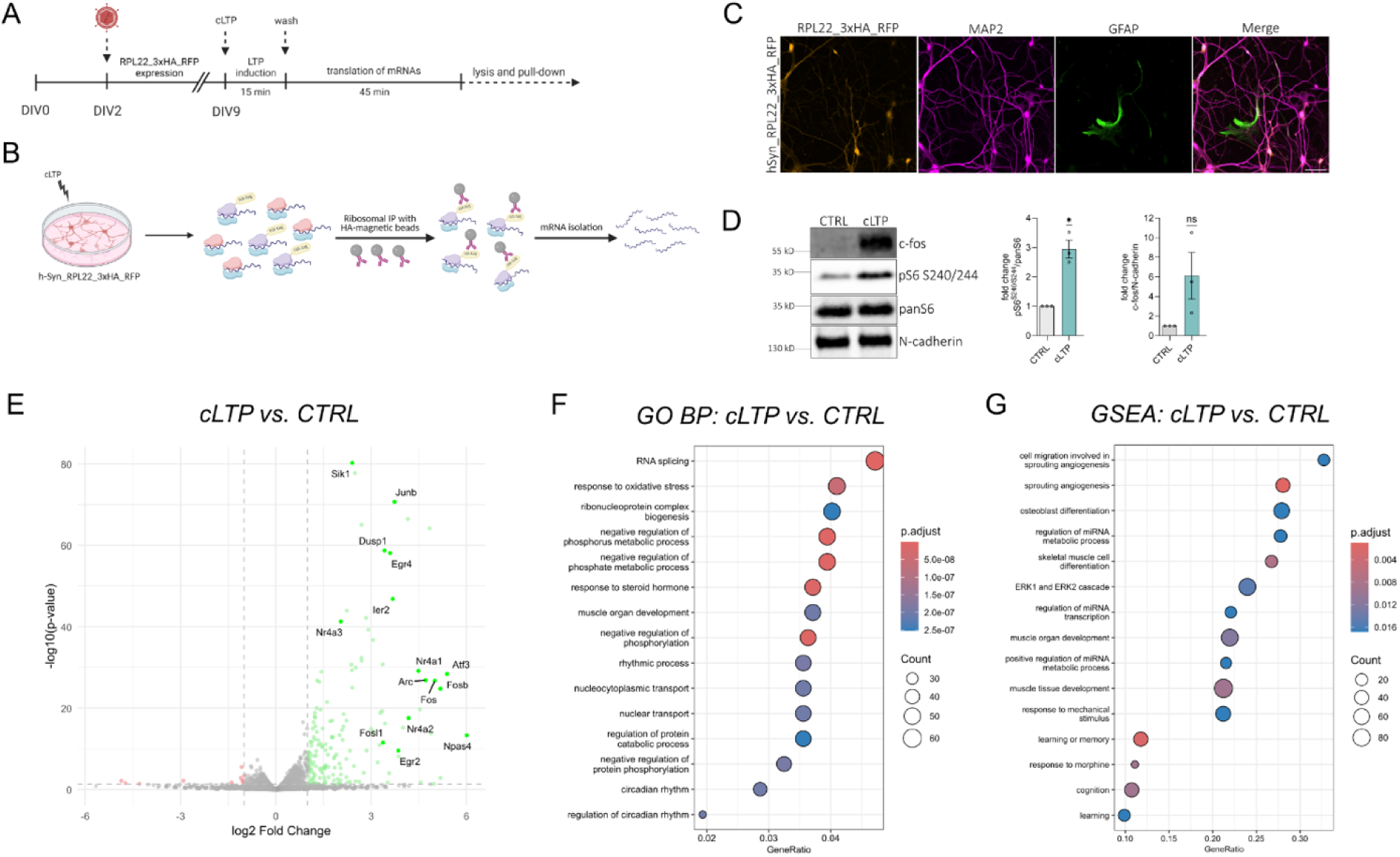
Translatome changes induced by neuronal stimulation in vitro. (**A**) Schematic of RiboTag experiment. Cultured cortico-hippocampal neurons were transduced with RPL22_3xHA_RFP expressing lentivirus on DIV 2. On DIV 9 neurons were stimulated for 15 min using a chemical long-term potentiation (cLTP) induction protocol. 1h post-stimulation cells were lysed and processed for RiboTag affinity chromatography and RNAseq analysis. (**B**) Graphical representation of newly translated proteins identified by RiboTag pull-down. Ribosomes containing RPL22_3xHA_RFP were immunoprecipitated by HA-magnetic beads. Co-precipitated mRNAs were isolated and analyzed by RNAseq. (**C**) Immunofluorescence images confirming neuron-specific expression of RPL22_3xHA_RFP. Red, RPL22_3xHA_RFP; green, GFAP, magenta, MAP2 to mark neuronal somata and dendrites. Scale bar, 50 µm. (**D**) Immunoblot showing mTORC1 signaling pathway activation in response to cLTP stimulation. Phosphorylation of ribosomal S6 increases upon cLTP. Right: mean (±SEM) change of S6 (S240/244) phosphorylation in control or cLTP neurons. n = 3 independent experiments compared by one sample t test. (**E**) Upregulation of immediate early gene (IEG) expression by cLTP induction of cortico-hippocampal neurons in vitro. Volcano plot showing expression changes induced by cLTP. Highlighted are upregulated (green) and downregulated (red) genes. Cut off: p-value < 0.05, log2FC > |1|. (**F**) Gene Ontology enrichment analysis of differentially translated proteins following cLTP stimulation. Circle size represents gene count, and colour indicates statistical significance (p-adjust). (**G**) Gene set enrichment analysis (GSEA) of differentially translated proteins following cLTP stimulation. (FDR < 0.05). Circle size represents gene count, and color indicates statistical significance (p-adjust).

Collectively, these data suggest that cLTP may serve as a starting point for the selection of candidate genes and pathways involved in activity-dependent memory consolidation.

To pursue this hypothesis further we applied our RiboTag and RNAseq approach to identify alterations in the layer 5b-specific translatome following exposure to an enriched environment (EE), a natural stimulation paradigm that evokes dendritic responses comparable to µStim.

### Enriched environment exposure increases IEG expression throughout the cortex

Exposure to an enriched-environment (EE) is a powerful approach to stimulate brain activity. To examine how a brief period of EE stimulation (30 min) influences brain plasticity, we applied a spatial transcriptomic approach to characterize region– and layer-specific transcriptional responses. Deconvolution analysis of EE datasets revealed a broad upregulation of immediate early genes (IEGs) across multiple cortical layers (Supp Fig. 3A–C). Notably, deep-layer excitatory neurons, showed the greatest number of upregulated IEGs (n = 12) relative to standard-environment controls. Several activity-regulated genes, including Fos and Sgk1, displayed distinct spatial expression patterns across brain regions (Supp Fig. 3D–H). Moreover, comparison of the EE spatial transcriptomic dataset with RiboTag cLTP data identified a significant overlap in differentially expressed IEGs (n = 6) (Supp Fig. 3I). Together, these findings indicate that even 30 minutes of EE robustly induces a defined set of IEGs, many of which are known to support experience-dependent plasticity and learning.

### Enriched environment exposure triggers translational remodeling in layer 5b neurons in vivo

Next we asked what proteins undergo translation in layer 5b pyramidal neurons of mice that receive sensory and cognitive experience in EE (Fig. 2 A–B). For this we used RiboTag mice expressing a ribosomal protein L10a fused to the GFP in the layer 5b neurons. L10a-GFP immunoprecipitation and RNA sequencing of translating mRNAs from somatosensory cortex revealed a marked increase in ribosome-associated transcripts relative to standard-housed controls (Fig. 2 C–E). Differentially expressed genes showed a significant overlap with those identified in the cLTP dataset, including canonical IEGs and regulators of cytoskeletal and synaptic remodeling (Fig. S1B).

**Figure 2.**
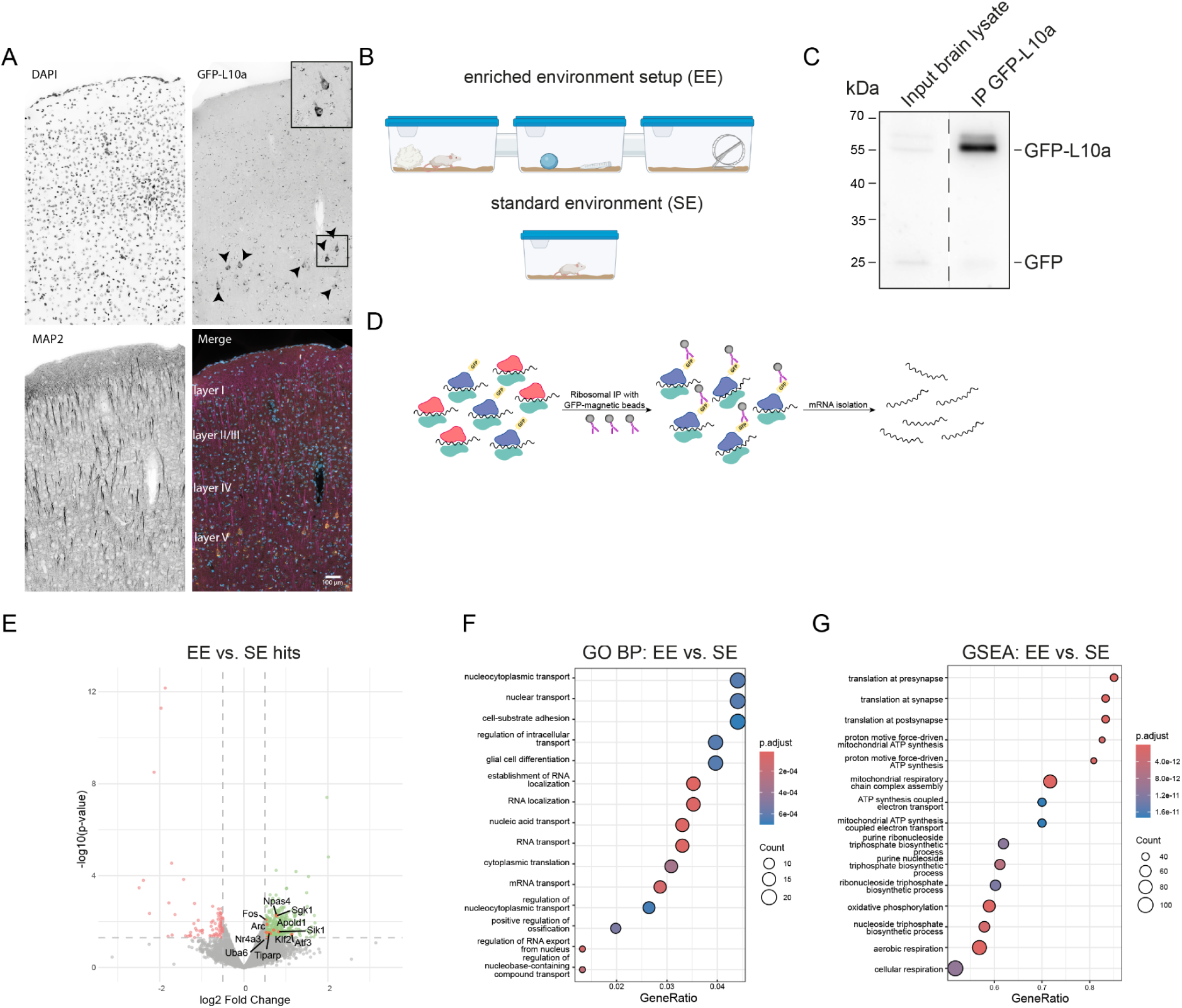
Layer 5b specific translatome after enriched environment exposure. (**A**) Specific expression of GFP-L10a ribosomal fusion protein in layer 5b of the somatosensory neocortex, enabling layer-specific RiboTag pull-down. Scale bar, 100 µm (**B**) Schematic of enriched environment (EE) exposure, providing enhanced sensory, motor and cognitive stimulation compared to standard housing (SE). Created with BioRender.com (**C**) Western blot of RiboTag pull-down from cortical homogenates, showing input and immunoprecipitated (IP) fractions. (**D**) Schematic of the RiboTag approach: genetically tagged ribosomal subunits are immunoprecipitated together with translating mRNAs, which are subsequently sequenced and analysed. (**E**) EE exposure upregulates translation of immediate early genes (IEGs). Volcano plot shows differentially expressed genes (DEGs; p < 0.05, log2FC > |0.5|), with highlighted overlap with cLTP datasets. (**F**) Gene Ontology (GO) analysis of biological processes reveals upregulation of genes related to mRNA transport and localization. (**G**) Gene Set Enrichment Analysis (GSEA) demonstrates an upregulation of genes involved in synaptic translation following EE exposure.

**Figure 3.**
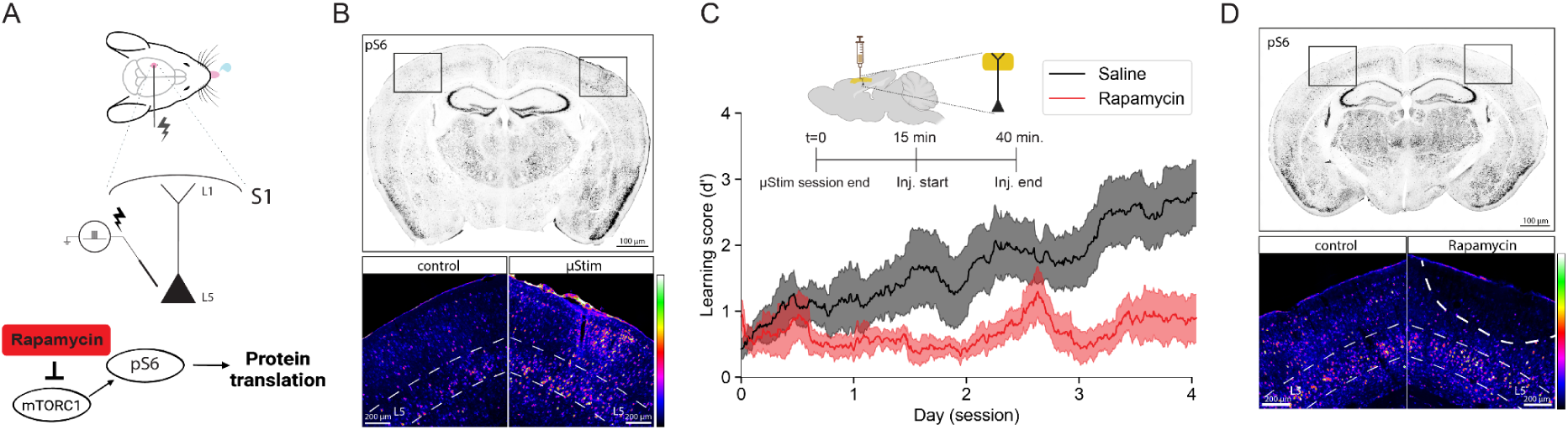
Microstimulation learning is mTORC1 dependent. (**A**) Schematic of the µStimulation learning paradigm, in which medial temporal lobe input to layer 1 tuft dendrites is necessary for learning the microstimulus-reward association and (below) a simplified representation of rapamycin action on the mTORC1 signaling pathway. (**B**) Representative pS6 immunohistochemistry in the stimulated and contralateral hemispheres, showing mTORC1-dependent activity. Scale bar overview: 100 µm; Zoom in: 200 µm (**C**) Behavioral learning curves for mice injected intracortically with saline or rapamycin immediately after each µStim session at 250 µm depth (nSaline=7, nRapamycin=5 mice); mTORC1 inhibition by rapamycin impairs or delays learning. (**D**) pS6 staining in control and rapamycin-injected hemispheres, illustrating reduced mTORC1 activity and the spatial spread of rapamycin. Scale bar overview: 100 µm; Zoom in: 200 µm.

Gene ontology and pathway analyses indicated enrichment for terms related to ‘mRNA transport’ and ‘RNA localization’, while gene-set enrichment analysis highlighted upregulation of translation at the synapse and postsynapse (Fig. 2 F–G).

Together, these findings show that environmental experience induces translation of a defined set of mRNAs in layer 5b neurons, consistent with an mTORC1-regulated dendritic plasticity program engaged during learning.

### Learning in the µStimulation paradigm requires mTORC1-dependent signaling in layer 1

We next tested whether mTORC1-regulated signaling in the neocortex is required for the consolidation of learning within the microstimulation (µStim) paradigm. In this task, head-fixed mice (Fig. 3A) were trained to associate intracortical electrical microstimulation targeting layer 5 (L5) of the primary somatosensory cortex (S1) with a timed water reward, an association that is known to depend on medial temporal lobe (MTL) input to cortical layer 1. Successful learning was quantified as an increase in sensitivity index (d’) (see Methods).

To validate the involvement of mTORC1 pathway activation, we assigned mice to two conditions stimControl (reward delivered independently of microstimulation) and stimLearn (reward dependent on microstimulation) (Fig S4). Immunohistochemical analysis revealed that microstimulation training was accompanied by increased phosphorylation of S6 ribosomal protein (pS6), a marker of mTORC1 pathway activation, in superficial cortical layers (L1, L2/3) (Fig. 3B).

To test the involvement of the mTORC1-pathway causality, we locally inhibited mTORC1 activity by infusing rapamycin or vehicle (saline) directly into superficial cortical layers (250µm depth), immediately after each training session. Immunostaining against pS6 indicated that mTORC1 was specifically inhibited in the upper cortical layers where apical and tuft dendrites of layer 5b neurons are located whereas the somatic mTORC1 was not affected (Fig. 3D). We then tested memory formation for four consecutive days. Inhibition of mTORC1 markedly reduced learning performance, reflected by decreased sensitivity index (d′) across sessions compared to controls, for all four days of learning (Fig. 3C) (Saline₍d₄₎:2.76 ± 0.50 vs. Rapamycin₍d₄₎:0.92 ± 0.36; Mann-Whitney U: *U*=37.0, *p*=0.022). This behavioral impairment was specific to post-training mTORC1 blockade and was not attributable to motor or motivational deficits.

These findings establish that mTORC1-dependent mechanisms in superficial cortex area necessary for memory consolidation in this cortical learning paradigm, linking synaptic input from hippocampal-temporal circuits to local mTORC1 signalling in the superficial layers of the cortex.

## Discussion

In this study we employed a complementary approach to define the translational landscape underlying cortical memory consolidation. We first utilized cLTP stimulation in cultured forebrain neurons to establish a broad candidate list of activity-dependent, ribosome-associated transcripts. By intersecting this broad dataset with layer 5b-specific RiboTag profiling following enriched environment (EE) exposure *in vivo*, we identified a robust translational signature intrinsic to layer 5b pyramidal neurons. This signature comprises immediate-early genes, synaptic regulators, and notably, key components of ribosome biogenesis and RNA handling. The specific upregulation of translational machinery in layer 5b neurons aligns with recent transcriptomic evidence showing an enrichment of ribosomal proteins in the excitatory synapses of cortical layer 1 (Spanò et al., 2025).

While persistent synaptic plasticity is thought to require *de novo* protein synthesis (Costa-Mattioli et al., 2009; Buffington et al., 2014), we further demonstrate via intracortical µStimulation that mTORC1 signalling is required for the consolidation of associative memory. mTORC1 integrates synaptic input through canonical pathways such as PI3K/Akt, enabling cap-dependent translation of plasticity-related mRNAs as well as metabolic regulation of protein turnover. This regulatory mechanism is a core element of long-term memory formation in cortical and hippocampal circuits (Buffington et al., 2014; Takei and Nawa, 2014). Our behavioural data, showing that post-training rapamycin impairs consolidation, directly support the notion that mTORC1 signalling in superficial cortical layers is required to stabilise learning-induced changes.

The molecular signatures of experience-driven translation and the underlying cell-type specificity identified in the RiboTag datasets show upregulation of candidate immediate-early genes (c-Fos, Fosl2, Arc), cytoskeletal regulators, RNA-binding proteins, and components of the local translation machinery. These signatures align with a broad body of work identifying local translation as a core cell-biological unit of neuronal computation (Holt and Schuman, 2013). Dendritic compartments rely on decentralised protein synthesis to support synapse-and dendritic compartment-specific plasticity within the relatively slow transport constraints of the soma. Additionally, our results sit well within modern frameworks demonstrating transcriptomic divergence across neuronal classes. For example, pyramidal neurons in different cortical layers display distinct transcriptional architectures (Grange et al., 2014), while interneuron subtypes are delineated by synaptic communication gene families (Paul et al., 2017). These findings suggest that the translational response we observe in layer 5b neurons may reflect a cell-type-specific molecular program adapted to the apical dendrite’s computational role.

Emerging evidence suggests that dendritic lysosomes may play a critical role in regulating local translation through their functional link to mTORC1 signalling (Khamsing et al., 2021). In our study, we observed up-regulation of translation initiation factors and ribosome-associated transcripts in layer 5b neurons, a finding that aligns with recent data showing an enrichment of ribosomal protein transcripts in excitatory synapses purified from layer 1 (Spanò et al., 2025). Although in our dataset we cannot discriminate between somatic and dendritic origin of mRNAs undergoing translation, it is possible that these transcripts are present in the distal dendrites. This arrangement implies that translation isn’t simply somatic but may be locally gated by organelle positioning and nutrient signalling state. Previous work suggests that actin-rich patches at dendritic shaft excitatory synapses cause stalling of actively transported lysosomes and therefore regulate lysosome positioning in the proximity of the synapse (Konietzny et al., 2017; Van Bommel et al., 2019). Such localised lysosomal stalling could provide a mechanism for spatially restricted metabolic signalling. Parallel studies show that lysosomes are recruited in an activity-dependent manner into dendrites and even to dendritic spines where they support synaptic protein turnover (Goo et al., 2017). Further, a recent review highlights that local translation, membranous organelle trafficking (including lysosomes) and synaptic plasticity are coordinated (Rajgor et al., 2021).

Importantly, activation of the Ragulator-lysosome complex relocates mTORC1 to late-endocytic/lysosomal compartments in neurons in response to BDNF/NMDA receptor stimulation (Khamsing et al., 2021). Although we did not directly image lysosomal dynamics in this study, the co-occurrence of translational machinery up-regulation and enrichment of mTORC1-regulated transcripts in dendritic tuft compartments is consistent with a model where local translation is coordinated by organelle positioning. Future work will need to determine if the dendritic lysosome acts as both a signalling checkpoint and local resource hub, linking distal synaptic activity to translational output necessary for memory consolidation.

A key mechanistic insight is that mTORC1 activation is spatially enriched in superficial layers, where the tuft dendrites of layer 5b pyramidal neurons reside. This supports a model in which dendritic compartments integrate memory-relevant long-range inputs from the medial temporal lobe and higher-order thalamus (Rubio-Garrido et al., 2009; Doron et al., 2020; Shin et al., 2021). This interpretation is consistent with work showing that distal apical dendrites express distinctive nonlinear and plasticity mechanisms. Thin apical dendrites possess specialised electrogenic properties, including NMDA spikes that act as high-gain amplifiers of converging distal inputs (Major et al., 2013). Beyond NMDA spikes, several forms of plasticity have been described in this region of the dendritic tree: spike-timing-dependent plasticity shaped by local conductances (Sjöström and Häusser, 2006), branch-specific calcium spikes and branch-level plasticity rules (Cichon and Gan, 2015), clustered-input supralinear events (Sandler et al., 2016), and regenerative dendritic events underlying compartmentalised computation (Otor et al., 2022; Benezra et al., 2024). Importantly, recent *in vivo* work has shown that apical dendrites undergo structural and functional plasticity shaped by behaviourally relevant input patterns, further reinforcing their role as dynamic integration sites that support learning (Pagès et al., 2021; Maristany De Las Casas et al., 2025).

We hypothesize that memory consolidation in the neocortex depends on spatially restricted molecular pathways operating at the apical tufts of layer 5b pyramidal neurons. Here, by positioning mTORC1 signalling and local translation machinery in the same compartment where long-range inputs converge, the cortex is equipped with a mechanism that can convert transient synaptic events into stabilised representations with branch-level precision. This framework offers a concrete molecular route by which systems-level information arriving in layer 1 is integrated, stored, and maintained over time, and highlights mTORC1-dependent signaling as a tractable target for understanding, and potentially modulating, cortical memory formation.

## Supplementary figures

**Fig. S1.**
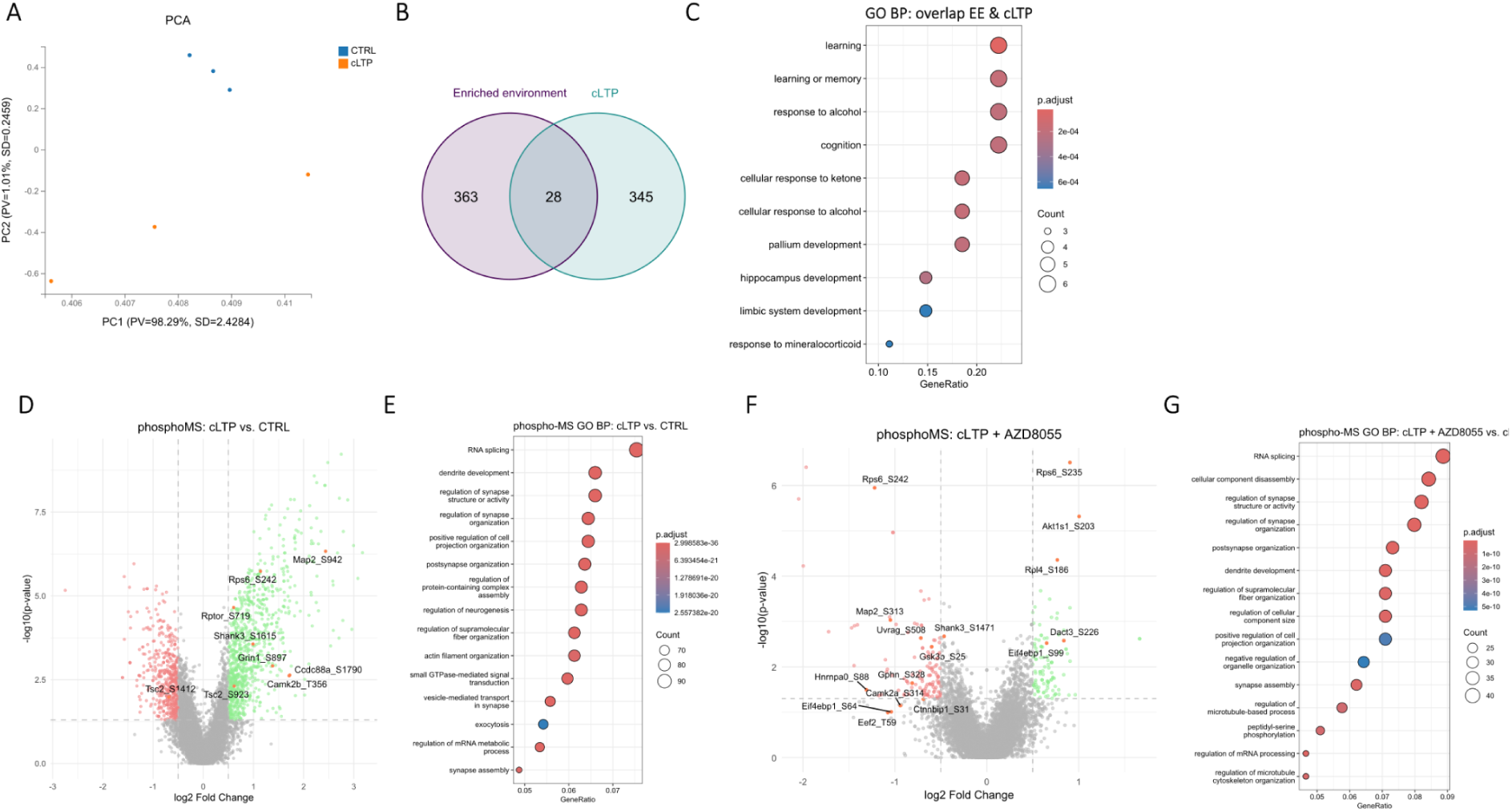
Translatome and phosphoproteome changes induced by neuronal stimulation in vitro. (**A**) Principal component analysis (PCA) of normalized RNA-seq counts from in vitro RiboTag experiment. Each point represents a biological replicate. (**B**) Venn diagram of genes which translation was upregulated in response to enriched environment (EE) in vivo and cLTP in vitro. (**C**) Gene Ontology enrichment analysis of genes which translation was upregulated in response to enriched environment (EE) in vivo and cLTP in vitro (see Figure S1. B). Bubble size represents gene count, and colour indicates statistical significance (p-adjust). (**D**) Stimulation of cortico-hippocampal neurons in vitro with cLTP protocol significantly upregulates phosphorylation of mTOR signalling components as well as components of postsynaptic density and cytoskeleton. Volcano plot showing phosphorylation changes induced by cLTP. Highlighted are upregulated (green) and downregulated (red) phosphosites. Cut off: p-value < 0.05, log2FC > |0.5|. (**E**) Gene Ontology enrichment analysis of differentially phosphorylated proteins following cLTP stimulation. Bubble size represents gene count, and colour indicates statistical significance (p-adjust). (**F**) Volcano plot showing phosphorylation changes in vitro during cLTP stimulation in the presence of mTOR kinase inhibitor AZD8055. Highlighted are upregulated (green) and downregulated (red) phosphosites. Cut off: p-value < 0.05, log2FC > |0.5|. (**G**) Gene Ontology enrichment analysis of differentially phosphorylated proteins following cLTP stimulation in the presence of mTOR kinase inhibitor AZD8055. Bubble size represents gene count, and colour indicates statistical significance (p-adjust).

**Fig. S2.**
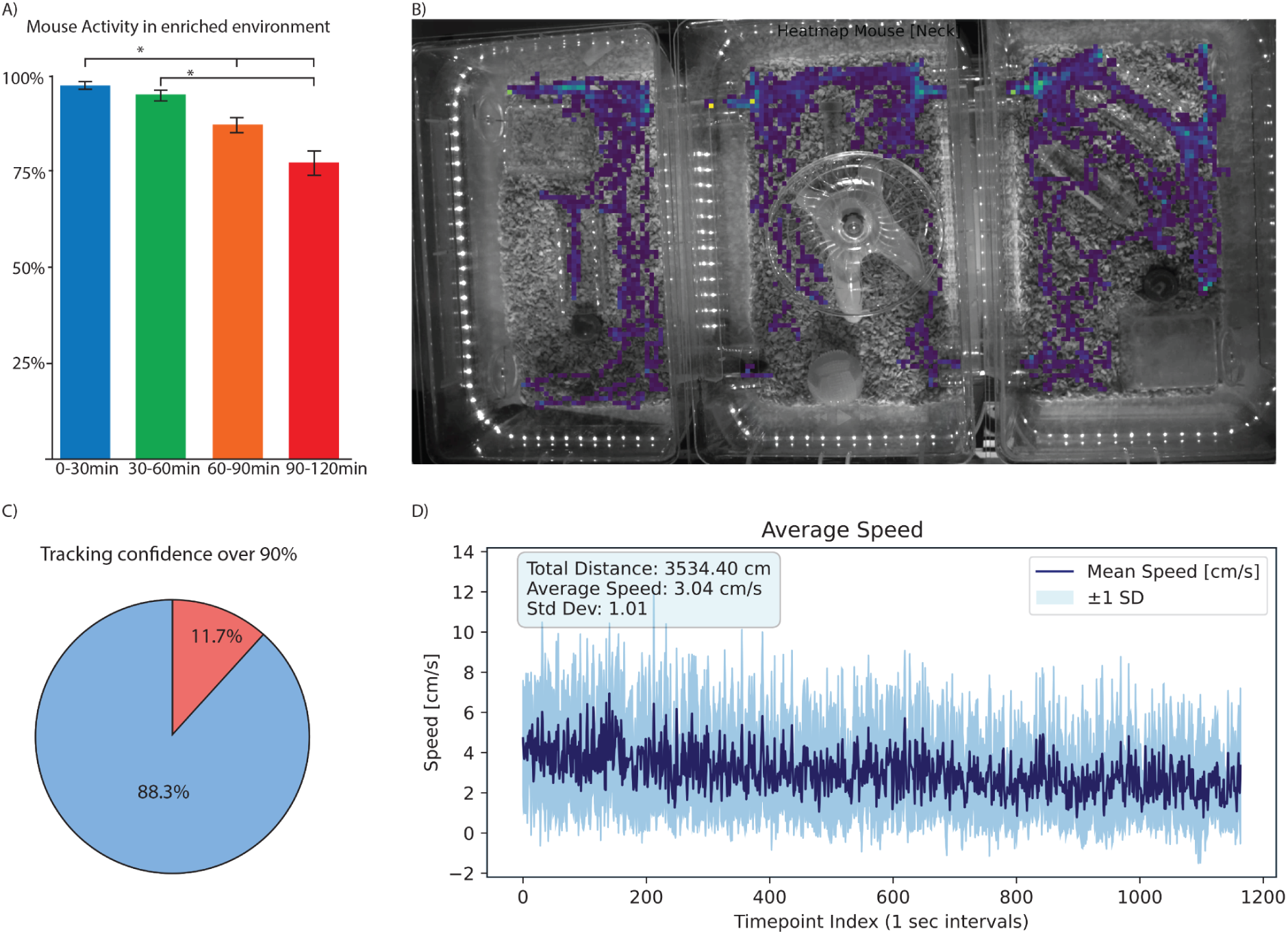
Automated tracking and activity analysis with ActiveMouse. (**A**) Exploratory activity of mice placed in an enriched environment or standard home cage, shown as the percentage of time spent exploring over a 120-min session. (**B**) Heat map generated by *ActiveMouse* illustrating the spatial distribution of mouse activity and frequently visited areas. (**C**) Tracking plot showing the total duration and trajectory of the mouse detected by *ActiveMouse*. (**D**) Summary plot of total distance travelled and movement speed. *ActiveMouse* integrates the DeepLabCut-Live framework with custom in-house code to enable automated, real-time monitoring and analysis of mouse behaviour without researcher presence.

**Fig. S3.**
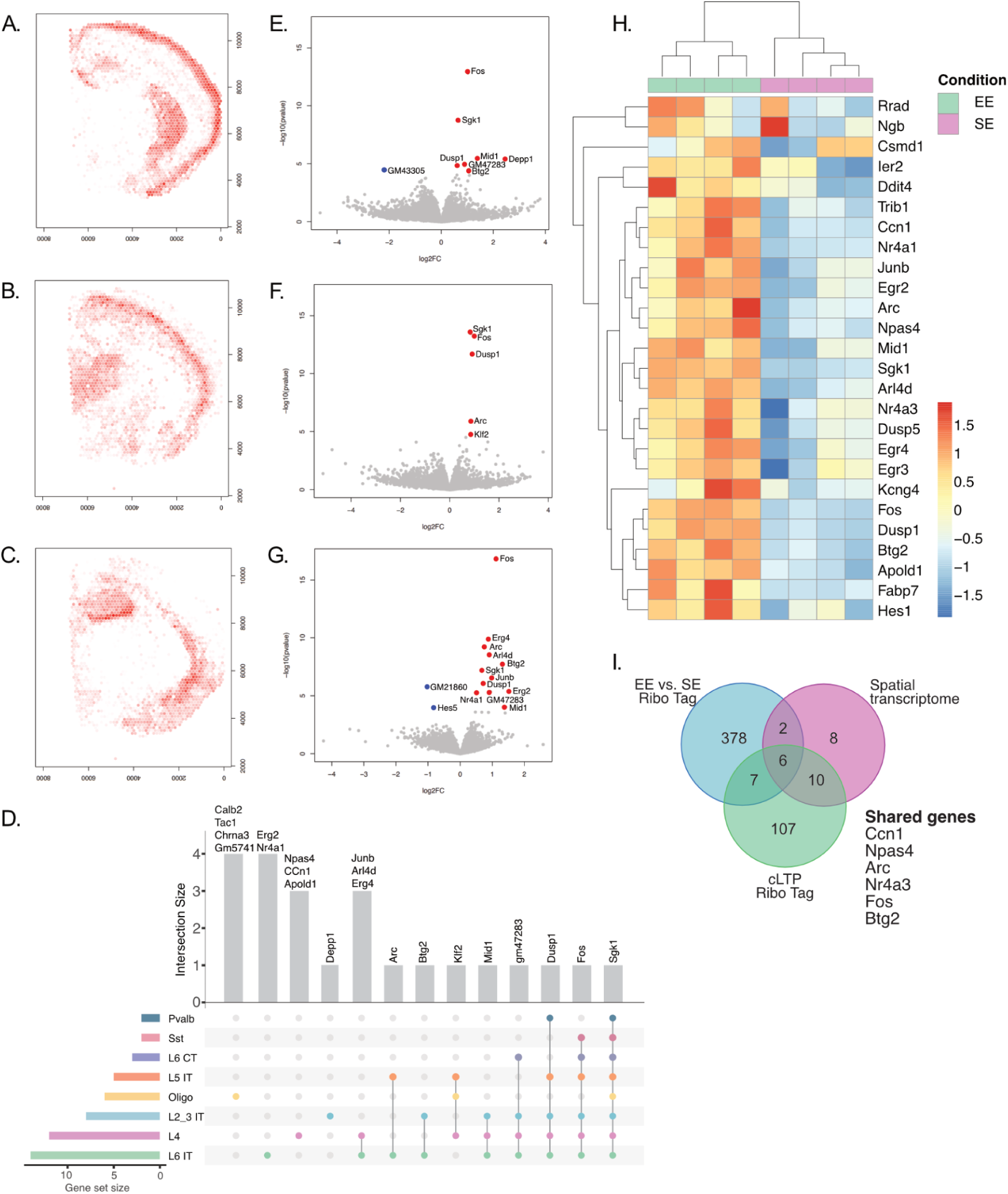
Spatial transcriptome after 30 minutes of enriched environment exposure. Deconvolution of spatial transcriptomic data using Allen Brain Institute as a reference for specific cell types. Red intensity shows expression of marker genes for the given cell types per spot **(A)** L2/3 IT – Layer 2/3 intra-telencephalic-projecting glutamatergic neurons **(B)** L5 IT – Layer 5 intra-telencephalic-projecting glutamatergic neurons **(C)** L6 IT – Layer 6 excitatory intra-cortical projection neurons. **(D)** Upset plot showing shared DEG each cell type when exposed to EE. Volcano plots showing DEG (DEG; P <0.05, log2FC > 0.05). Red is up regulated and blue is down regulated after EE exposure for the following cell types: **(E)** Layer 2/3 intra-telencephalic-projecting glutamatergic neurons, **(F)** Layer 5 intra-telencephalic-projecting glutamatergic neurons, **(G)** Layer 6 excitatory intra-cortical projection neurons. **(H)** Heat map showing gene expression of IEG between EE and SE across four brain hemispheres. **(I)** Venn diagram showing shared DEG across EE vs. SE Ribo Tag experiment, cLTP Ribo Tag experiments and EE vs. SE spatial transcriptomic experiments.

**Fig. S4.**
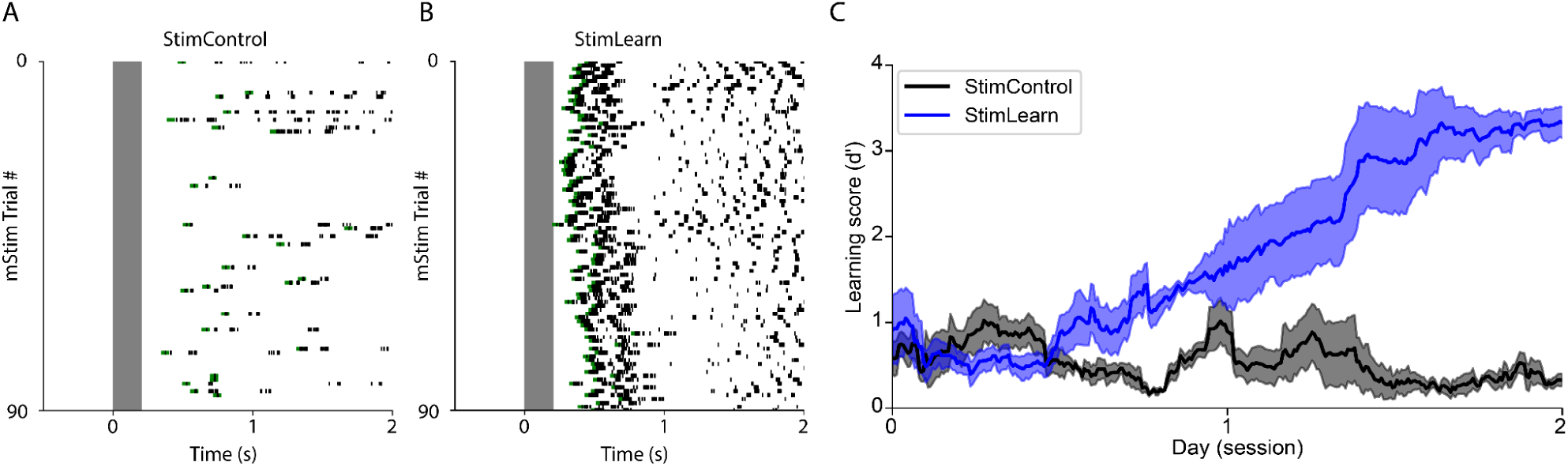
Behaviour of animals for immunohistochemistry. **(A)** Example licking behaviour for all microstimulation trials of a mouse on the final day of microstimulation training for a StimControl group. Green dots indicate the reward lick, black dots correspond to consecutive licks. **(B)** Example behaviour of a mouse on the final day of microstimulation training for a StimLearn group. **(C)** Learning score over 2 days comparing StimControl to StimLearn groups. (nStimControl=4, nStimLearn=4 mice, StimControl₍t_300_₎:0.032±0.06 vs. StimLearn₍t_300_):3.29±0.20; Independent T-test: t=-12.54, p<0.01; data presented as mean ± SEM).

## Notes

### Competing Interest Statement

The authors have declared no competing interest.

### Funding

- Deutsche Forschungsgemeinschaft: 327654276 (CRC1315, subproject A10)

