## Supplemental figures for "mTORC1-Dependent Signaling in Layer 5b Neurons Is Required for Memory Consolidation"

### Supplementary figures

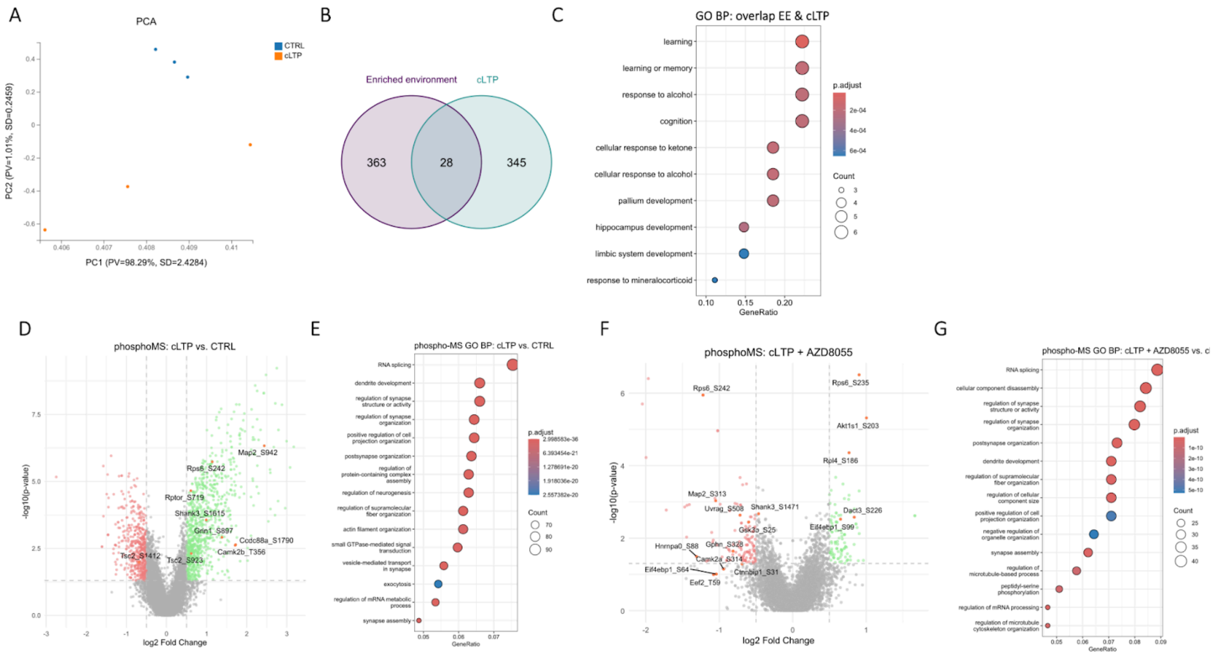


*Fig. S1. Translatome and phosphoproteome changes induced by neuronal stimulation in vitro.*

(**A**) Principal component analysis (PCA) of normalized RNA-seq counts from in vitro RiboTag experiment. Each point represents a biological replicate. (**B**) Venn diagram of genes which translation was upregulated in response to enriched environment (EE) in vivo and cLTP in vitro. (**C**) Gene Ontology enrichment analysis of genes which translation was upregulated in response to enriched environment (EE) in vivo and cLTP in vitro (see Figure S1. B). Bubble size represents gene count, and colour indicates statistical significance (p-adjust). (**D**) Stimulation of cortico-hippocampal neurons in vitro with cLTP protocol significantly upregulates phosphorylation of mTOR signalling components as well as components of postsynaptic density and cytoskeleton. Volcano plot showing phosphorylation changes induced by cLTP. Highlighted are upregulated (green) and downregulated (red) phosphosites. Cut off: p-value < 0.05, log2FC > |0.5|. (**E**) Gene Ontology enrichment analysis of differentially phosphorylated proteins following cLTP stimulation. Bubble size represents gene count, and colour indicates statistical significance (p-adjust). (**F**) Volcano plot showing phosphorylation changes in vitro during cLTP stimulation in the presence of mTOR kinase inhibitor AZD8055. Highlighted are upregulated (green) and downregulated (red) phosphosites. Cut off: p-value < 0.05, log2FC > |0.5|. (**G**) Gene Ontology enrichment analysis of differentially phosphorylated proteins following cLTP stimulation in the presence of mTOR kinase inhibitor AZD8055. Bubble size represents gene count, and colour indicates statistical significance (p-adjust).

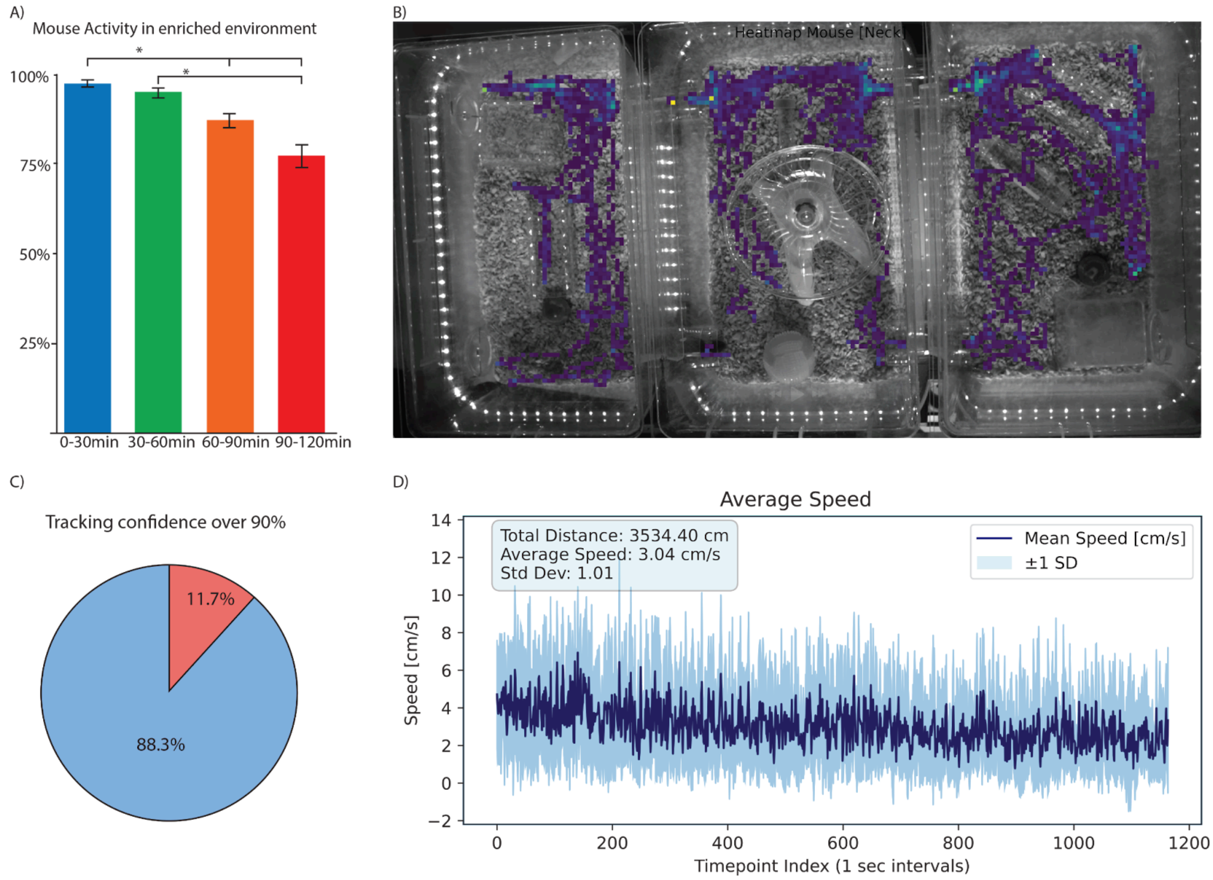


*Fig. S2 Automated tracking and activity analysis with ActiveMouse.*

(**A**) Exploratory activity of mice placed in an enriched environment or standard home cage, shown as the percentage of time spent exploring over a 120-min session. (**B**) Heat map generated by *ActiveMouse* illustrating the spatial distribution of mouse activity and frequently visited areas. (**C**) Tracking plot showing the total duration and trajectory of the mouse detected by *ActiveMouse*. (**D**) Summary plot of total distance travelled and movement speed. *ActiveMouse* integrates the DeepLabCut-Live framework with custom in-house code to enable automated, real-time monitoring and analysis of mouse behaviour without researcher presence.

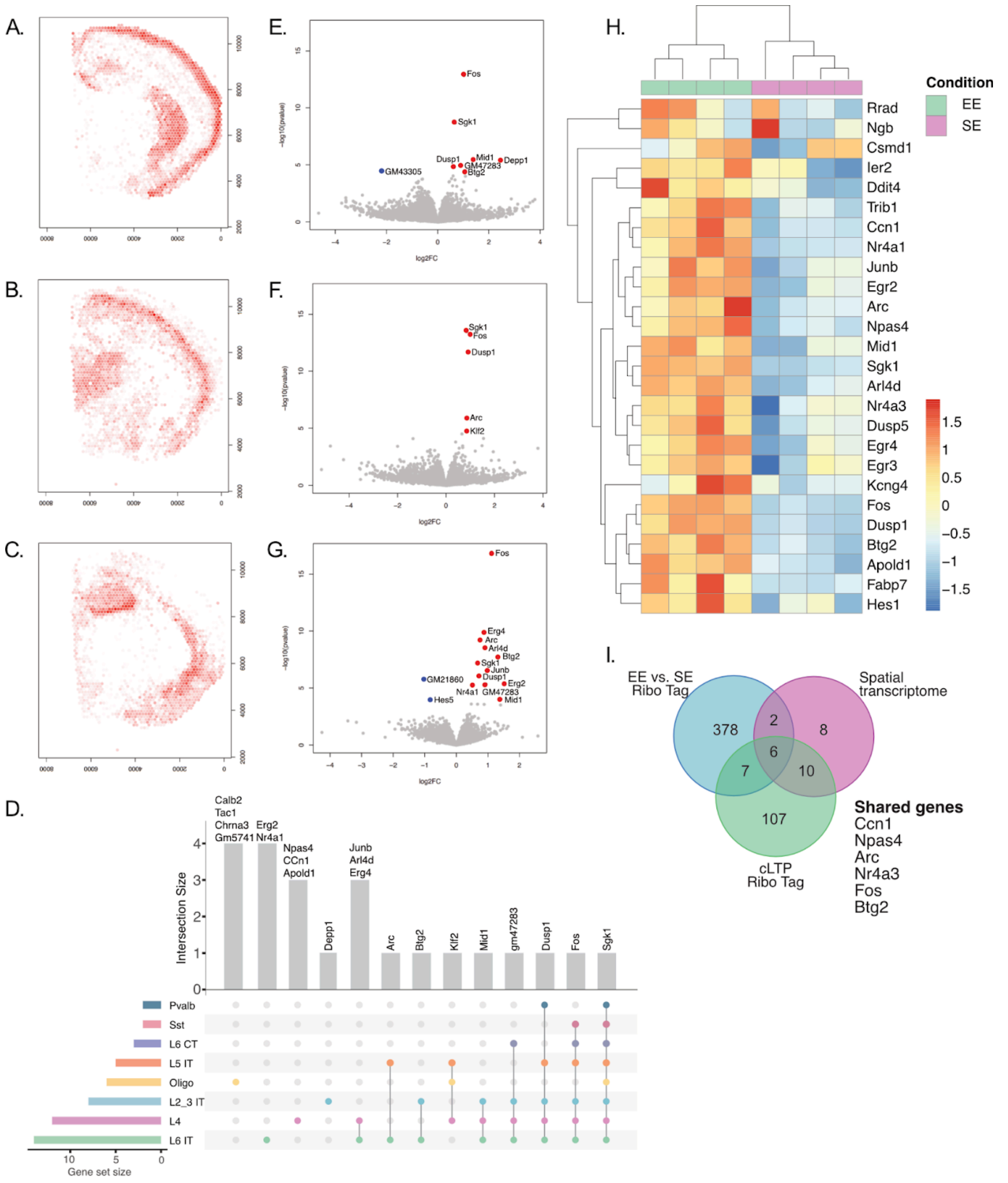


*Fig. S3. Spatial transcriptome after 30 minutes of enriched environment exposure*

Deconvolution of spatial transcriptomic data using Allen Brain Institute as a reference for specific cell types. Red intensity shows expression of marker genes for the given cell types per spot **(A)** L2/3 IT - Layer 2/3 intra-telencephalic-projecting glutamatergic neurons **(B)** L5 IT - Layer 5 intra-telencephalic-projecting glutamatergic neurons **(C)** L6 IT - Layer 6 excitatory intra-cortical projection neurons. **(D)** Upset plot showing shared DEG each cell type when exposed to EE. Volcano plots showing DEG (DEG; P <0.05, log2FC > 0.05). Red is up regulated and blue is down regulated after EE exposure for the following cell types: **(E)** Layer 2/3 intra-telencephalic-projecting glutamatergic neurons, **(F)** Layer 5 intra-telencephalic-projecting glutamatergic neurons, **(G)** Layer 6 excitatory intra-cortical projection neurons. **(H)** Heat map showing gene expression
of IEG between EE and SE across four brain hemispheres. **(I)** Venn diagram showing shared DEG across EE vs. SE Ribo Tag experiment, cLTP Ribo Tag experiments and EE vs. SE spatial transcriptomic experiments.

*Fig. S4 - Behaviour of pS6 immuno histochemistry*

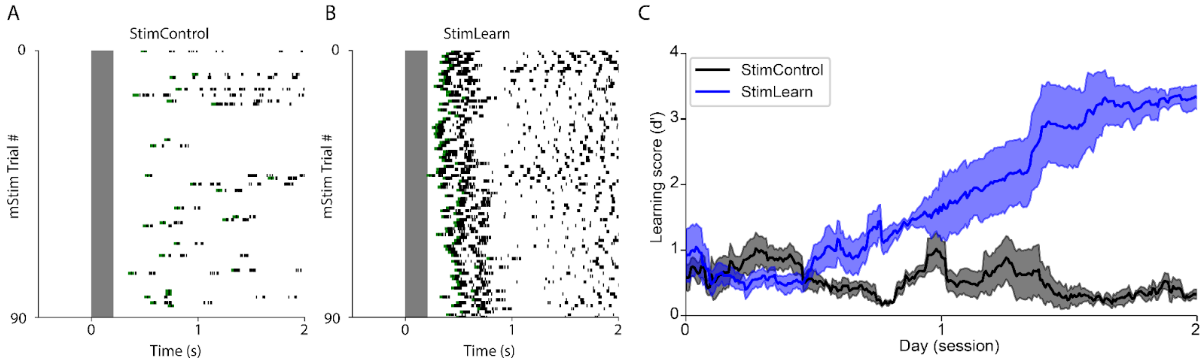


*Fig. S4. Behaviour of animals for immunohistochemistry.*

**(A)** Example licking behaviour for all microstimulation trials of a mouse on the final day of microstimulation training for a StimControl group. Green dots indicate the reward lick, black dots correspond to consecutive licks. **(B)** Example behaviour of a mouse on the final day of microstimulation training for a StimLearn group. **(C)** Learning score over 2 days comparing StimControl to StimLearn groups. (nStimControl=4, nStimLearn=4 mice, StimControl$(t_{300})$:0.032±0.06 vs. StimLearn$(t_{300})$:3.29±0.20; Independent T-test: t=-12.54, p<0.01; data presented as mean ± SEM).
